# Fungivorous protists in the rhizosphere of *Arabidopsis thaliana* – Diversity, functions, and publicly available cultures for experimental exploration

**DOI:** 10.1101/2023.07.12.548669

**Authors:** Antonie H. Estermann, Justin Teixeira Pereira Bassiaridis, Anne Loos, Marcel Dominik Solbach, Michael Bonkowski, Sebastian Hess, Kenneth Dumack

## Abstract

In the context of the soil food web, the transfer of plant-fixed energy and carbon to higher trophic levels has traditionally been attributed to two main energy channels: the fungal energy channel and the bacterial energy channel. Historically, protists were overlooked in the fungal energy channel, which was believed to be controlled by fungivorous microarthropods and nematodes. In this study, we investigated fungivorous protists in the rhizosphere of *Arabidopsis thaliana*. Our findings revealed a notable abundance and diversity of protists that have developed specialized strategies to overcome the protective cell wall of fungi. Among the identified species were two Vampyrellida (Rhizaria) species, namely *Theratromyxa weberi* and *Platyreta germanica*, as well as one Arcellinida (Amoebozoa) species, called *Cryptodifflugia oviformis*. While *T. weberi* typically consumed entire fungal cells, the other two species perforated fungal cell walls and extracted the cellular contents. We elucidate the feeding strategies and dietary ranges of the amoebae, highlighting the non-uniform nature of fungivory in protists, as different taxa have evolved distinct approaches to access fungi as a food source. Moreover, we provide publicly available cultures of these protists to facilitate further experimental investigations within the research community.

## 1. Introduction

In the soil food web, energy and carbon fluxes are traditionally considered to start at the plants, the photosynthetic primary producers. Subsequently, this plant-fixed carbon was expected to be channeled to higher trophic levels by two main energy channels, the fungal and bacterial energy channels (Moore et al., 1988; Holtkamp et al., 2011; Bradford, 2016). The bacterial energy channel was believed to rely on bacterivorous protists and nematodes. The fungal energy channel was traditionally seen to be controlled by fungivorous microarthropods and nematodes (Hunt et al., 1987; de Ruiter et al., 1995). However, the plant holobiont, i.e. the sum of the plant and its associated microorganisms, was found to be much more complex. These plant-associated microorganisms compete and interact with each other, form complex food webs, and their community composition is shaped by both top-down and bottom-up effects (Hartmann et al., 2009; Krashevska et al., 2016). The plant-related functions of microorganisms include nutrient acquisition, disease suppression, and the increase in stress tolerance (Berendsen et al., 2012; Richardson, 2017; Hassani et al., 2018; Xiong et al., 2020). Thus, microorganisms are essential for plant health.

Protists associated with soils and plants were long considered one-dimensional, simple bacterivores, likely because their majority has been isolated and maintained with bacteria as a food source (Bass et al., 2009b; Howe et al., 2009; Glücksman et al., 2010; Geisen et al., 2015a). However, there are numerous reports of soil protists that specifically interact with fungi and microscopic animals, and, in addition, some protists can process multiple food sources (bacteria, fungal cells, algae). Those taxa can be considered non-specialized, omnivorous phagotrophs, which are of high functional versatility (Dumack et al., 2019). Ekelund (1998) found that omnivorous protists feeding on fungal cells may reach similar biomass as bacterivorous protists. This suggests that they can have significant impact on soil food webs and plant health.

So far, taxonomic surveys report a relatively limited diversity of soil-dwelling protists that evolved special adaptions to feed on fungi, including their large hyphal structures (Petz et al., 1985; Ogden and Pitta, 1990; Pakzad, 2003). The most prominent examples belong to the ciliates (Grossglockneridae, Ciliophora), vampyrellid amoebae (Vampyrellida, Rhizaria), and testate lobose amoebae (Arcellinida, Amoebozoa). It takes specialist knowledge to isolate and cultivate such fungivorous protists, and due to the physical nature of soils and plant surfaces, extra efforts to study their poorly characterized interactions with other organisms.

Recent environmental sequencing studies indicate that certain protists shape the fungal community and best explain the suppression of certain fungal pathogens (Huang et al., 2021; Ren et al., 2023). However, the experimental exploration of the mechanisms that underlie these findings was hampered by the lack of microscopical observations and publicly available cultures. In this study, we screened roots of *Arabidopsis thaliana* from natural populations for fungivorous protists. We isolated and characterized several strains of more or less specialized, fungivorous amoebae, and provide their cultures to the scientific community to facilitate the experimental exploration of plant-fungi-protist relationships.

## 2. Material and methods

### 2.1. Sampling and isolation

Six *Arabidopsis thaliana* were sampled around Cologne, Germany. Roots were washed and cut into 1 cm long pieces. These root pieces were transferred into the wells of a 24-well plate with the culture medium Waris-H+Si (Mcfadden and Melkonian, 1986), and incubated for one day. The wells were screened with an inverted phase contrast microscope for fungivorous protists (Nikon Eclipse TS100; 100x - 400x). From the total of 144 wells, almost every well contained numerous *Cryptodifflugia* specimens and four wells with vampyrellid cells were found. Cultures were established with *Saccharomyces cerevisiae* as prey. Cell sorting as described in Solbach et al. (2021) or serial manual washings in sterile medium were used to establish cultures free of environmental bacteria. Microscopic documentation was done with a Nikon Eclipse 90i and Motic AE2000.

### 2.2. Molecular identification and phylogenetics

To identify the strains by molecular means, the SSU rRNA gene was amplified and sequenced. For the vampyrellid strains, digestive cysts were picked and transferred to 1 ml of fresh culture medium. To avoid any contamination by food organisms during PCR, the cysts were starved for several days until they had hatched. The resulting amoebae were then transferred with approx. 1 μl of medium to PCR tubes filled with 4 μl ddH_2_O and stored at -20 °C. For *Cryptodifflugia*, cells in 5 μl of medium were transferred into PCR tubes. The PCR mixture was added, consisting of 6.4 μl ddH2O, 1.7 μl Thermo Scientific Dream Taq Green Buffer, 1.7 μl of 10 μM forward and reverse primers each, 0.34 μl 10 μM dNTPs and 0.17 μl DreamTaq polymerase (Thermo Fisher Scientific, Dreieich, Germany). The SSU rRNA genes were amplified in two successive steps. First, the whole SSU rRNA gene was amplified with the general eukaryotic primers EukA and EukB (Medlin et al. 1988). In the second step, semi-nested re-amplifications were performed. For Vampyrellida, we used Vampyrellida-specific primers in combination with eukaryotic primers, i.e. EukA + 947_Vamp (5’-AAGAAGATATCCTTGGT-3’; Fiore-Donno et al., 2020) targeting the 5’-part of the SSU rDNA and 615_Phyt (5’-CTTTSAARGCTCGTAGTTG-3’; Fiore-Donno et al., 2020) + EukB for the 3’-part of the gene. One microlitre of the first PCR product was used as a template for reamplification. For *Cryptodifflugia*, the semi-nested re-amplifications were performed by combining EukA and EukB with Phryganellina-specific primers, i.e. 590F_Phry (5’-CGGTAATTCCAGCTCCAATAGT-3’; Dumack et al., 2020) and 1300R_Phry (5’-GCATGGCCGTTTTTAGTTGGTG-3’; Dumack et al., 2020) as described above. PCR conditions were as follows: initial denaturation at 95 °C for 5 min, 24 cycles (denaturation at 96 °C for 32 s, annealing at 50 °C for 36 s, elongation at 72 °C for 2 min), terminal extension at 72 °C for 7 min, and hold at 4 °C. The PCR products were purified by adding 0.15 μl of Exonuclease I, 0.9 μl FastAP and 1.95 μl water to 8 μl of the second PCR product and then heated for 30 min at 37 °C, and subsequently for 20 min at 85 °C. The Big dye Terminator Cycle sequencing Kit (Thermo Fisher Scientific, Dreieich, Germany) and an ABI PRISM automatic sequencer were used for the sequencing. All four previously mentioned primers were used for sequencing.

For the vampyrellid phylogeny, the SSU rRNA gene sequences were added to a sub-alignment (Leptophryidae with Vampyrellidae as outgroup) of that used in Hess and Suthaus (2022), and then aligned with MUSCLE in Seaview 5.0.4 (Gouy et al., 2010) and curated manually. We then inferred maximum likelihood (ML) and Bayesian (BI) phylogenies with 1,541 aligned sites. Twenty maximum likelihood inferences were done with raxmlGUI 2.0.10 (Edler et al., 2021) with the GTR+I+Γ model, and the best tree was selected for display. The branch support was retrieved by bootstrapping with 1,000 replicates. The Bayesian analysis was done with Beast 2.7.3. (Bouckaert et al., 2014) with the same model, 5,000,000 generations (trees sampled every 1,000 generations), and a 25 % burn-in (1,250,000 generations discarded). We confirmed the stationarity of the BI with Tracer 1.7.2. (Rambaut et al., 2018).

For the Arcellinida phylogeny placing *Cryptodifflugia*, the sequences were added to a sub-alignment of that used in Dumack et al. (2020), and then aligned with MUSCLE in Seaview 5.0.4 and curated manually. We then inferred maximum likelihood (ML) and Bayesian (BI) phylogenies with 1,321 aligned sites, which were 40.88 % invariant. We used the model GTR+I+Γ. For phylogenetic calculations, we used MrBayes v3.2.6 (Ronquist and Huelsenbeck, 2003) and RAxML (Stamatakis, 2014). The Bayesian analysis was set up with a sampling of every 100 and a diagnosis of every 500 trees and 25 % of burn-in. It was conducted on the online platform CIPRES (Miller et al., 2010) and reached final split frequencies of <0.01 after 870,000 generations.

### 2.3. Food range experiment

Twelve different fungal species were used as potential food sources for the amoebae and cultivated in Petri dishes on potato glucose agar (Sigma-Aldrich) according to the manufacturer’s instructions. For the vampyrellid amoebae, a small quantity of each fungal culture was given into half-strength Waris-H+Si and transferred to wells of a twelve-well plate. The wells were observed after one, seven, and fourteen days for growth of the protists and signs of predation with the AE2000 (Motic) inverted microscope and the MikroLive 6,4MP camera. For *Cryptodifflugia*, a small quantity of each fungal culture was transferred to separate Petri dishes containing 1% water agar. After a few days of incubation, about 300 μl of a fungi-free *Cryptodifflugia* culture in WG medium (Bonkowski, 2019) was added to each fungal subculture. The preparation was then filmed with the Nikon Eclipse TE2000 inverted microscope and the Nikon digital sight DS-U3 camera (program: NIS-Elements V4.13.04, Tokyo, Japan) after one, seven, and fourteen days for amoebal growth and signs of predation. Time-lapse videos were prepared to confirm predation on each fungal culture.

### 2.4. Phalloidin staining and fluorescence microscopy

To obtain first insights into the mechanism used by *Cryptodifflugia oviformis* to feed on fungi, cells were fixed and stained with a fluorescent phalloidin-conjugate before and during the feeding process. For this, gelatin-coated cover glasses were given into 6-well plates and sterilized with ultraviolet light for 20 minutes. Subsequently, WG medium with hyphae of *Amniculicola* sp. was pipetted onto the cover glasses. Excess medium around the hyphae was removed. When hyphae adhered to the cover glasses, a culture of *Cryptodifflugia oviformis* was added and the setup was incubated for up to 24 h. After incubation, the cells were fixed with 4 % formaldehyde (freshly made from paraformaldehyde dissolved in PBS, pH 7) for 10 min at room temperature. The coverslips were subsequently washed with PBS (Sigma Aldrich) for 5 min. After fixation, the cells were permeabilized for 10 min with 0.1 % Triton X-100 in TBS. Subsequently, slides were washed with PBS for 5 min and stained with Phalloidin-iFluor 488 (ab176753, abcam, 0.001 %). Incubation took place for 1 h in the dark (in a moist chamber, room temperature), followed by three washing steps (PBS, each 5 min). Afterwards, mounting medium (85 % glycerol, 15 % PBS) was added and the cells were documented with a Nikon Eclipse 90i.

#### Scanning electron microscopy

To obtain insights into the mechanism used by *Platyreta germanica* to feed on fungi, the supernatant of an old culture of *Platyreta germanica* strain G2.3 was carefully removed and the emptied yeast cells resuspended by shaking. The suspension was placed on poly-L-lysine coated coverslips and let settle for about one hour. The coverslips were then transferred to ethanol/water mixtures of ascending concentrations (30 %-50 %-70 %-90 %-100 %), and finally immersed twice in hexamethyldisilazane (HMDS) for at least 5 min. After careful removal of HMDS the coverslips were air-dried in a fume hood, sputter-coated with gold and imaged with a ZEISS Neon 40 scanning electron microscope using the secondary electron detector (ZEISS, Oberkochen, Germany).

## 3. Results

### Genetic identity of the isolated strains

Thirteen strains of naked and testate amoebae were isolated from the roots of *Arabidopsis thaliana* and cultivated with *Saccharomyces cerevisiae* as food source. Based on their SSU rRNA gene sequences, the isolates could be assigned to two distinct and distantly related orders of amoeboid life forms, the Vampyrellida (Rhizaria) and Arcellinida (Amoebozoa) (Figure 1A). Phylogenetic inferences revealed that the four vampyrellid isolates branched in the family Leptophryidae with maximum support (Figure 1B). Two strains (G7.2, U11) were closely related to *Theratromyxa weberi* (strain ATCC 50200), the other two (G2.3, S1P3) to *Platyreta germanica* (strains “Gross-Gerau” and “Fritzlar-Werkel”). As the genetic distances between the new strains and the published reference sequences were <0.5 %, we assign the new strains to these two species (Table 1). The nine arcellinid strains branched within the family Cryptodifflugiidae with maximum support (Figure 1C). They are all closely related to each other and to *Cryptodifflugia oviformis* (JQ366062) and comply with the morphological circumscription of this species (details below). The cultures of *Theratromyxa weberi* and *Platyreta germanica* could be grown without environmental bacteria, while bacteria-free cultures of *Cryptodifflugia oviformis* showed atypical behavior, i.e. daughter cells lacked shells and ceased to grow after approximately five divisions. Hence, the here presented cultures of *Cryptodifflugia oviformis* contained environmental bacteria throughout subsequent analyses.

**Table 1:**
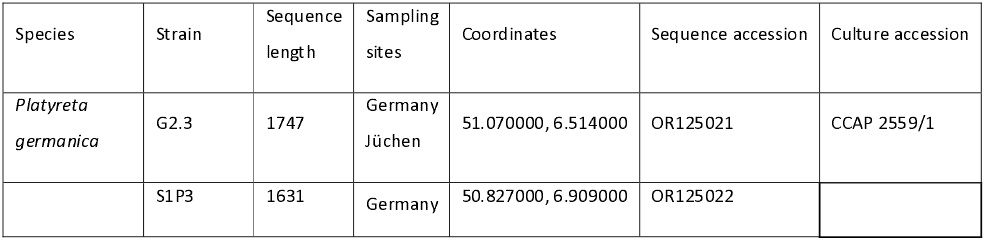
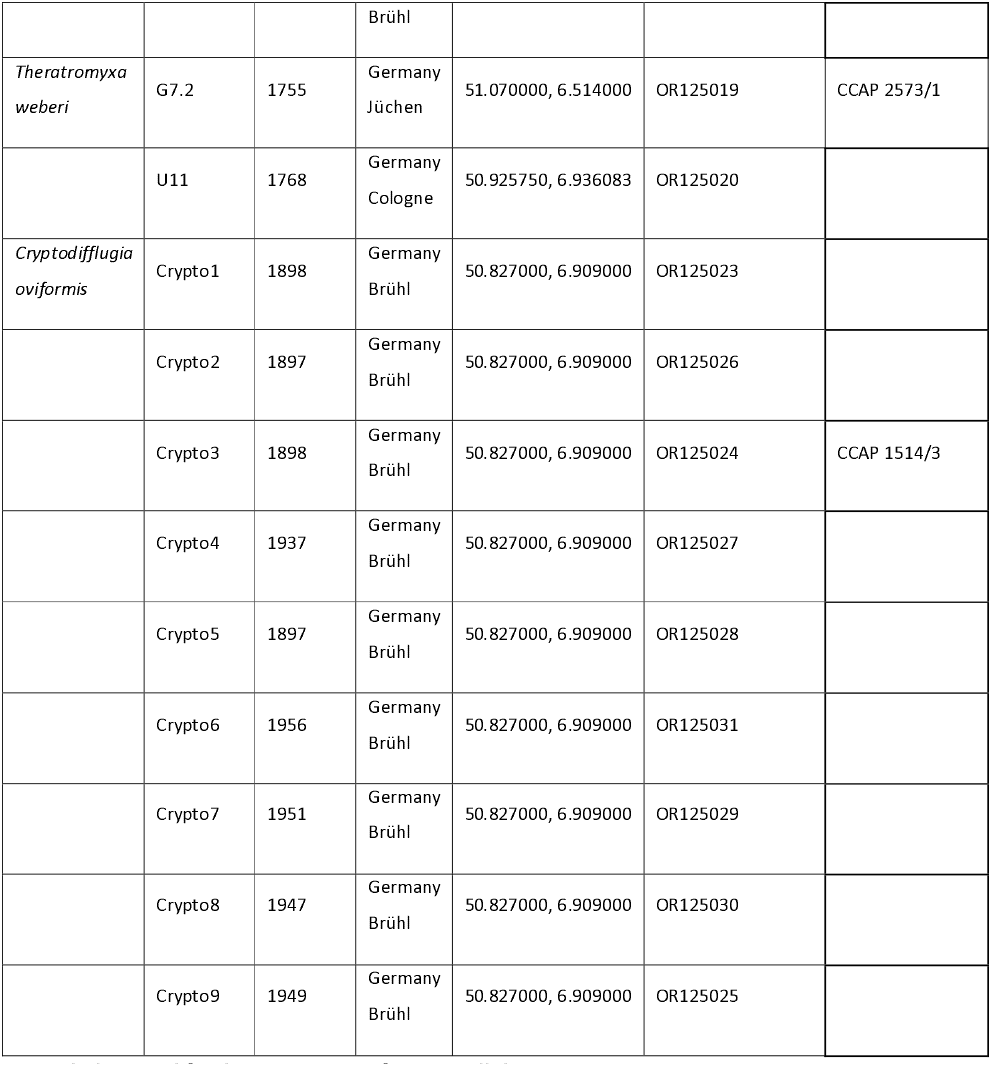
Investigated strains of fungivorous protists and corresponding data.

**Table 2:**
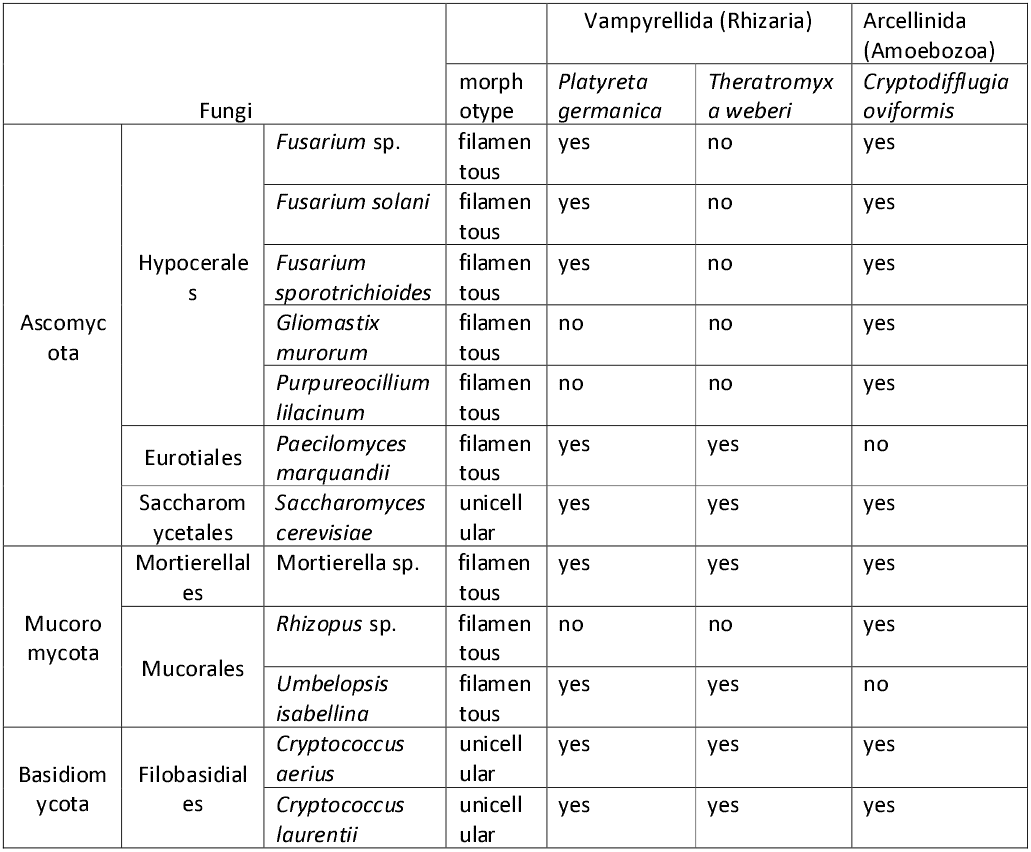
Results of the food range experiments.

**Figure 1:**
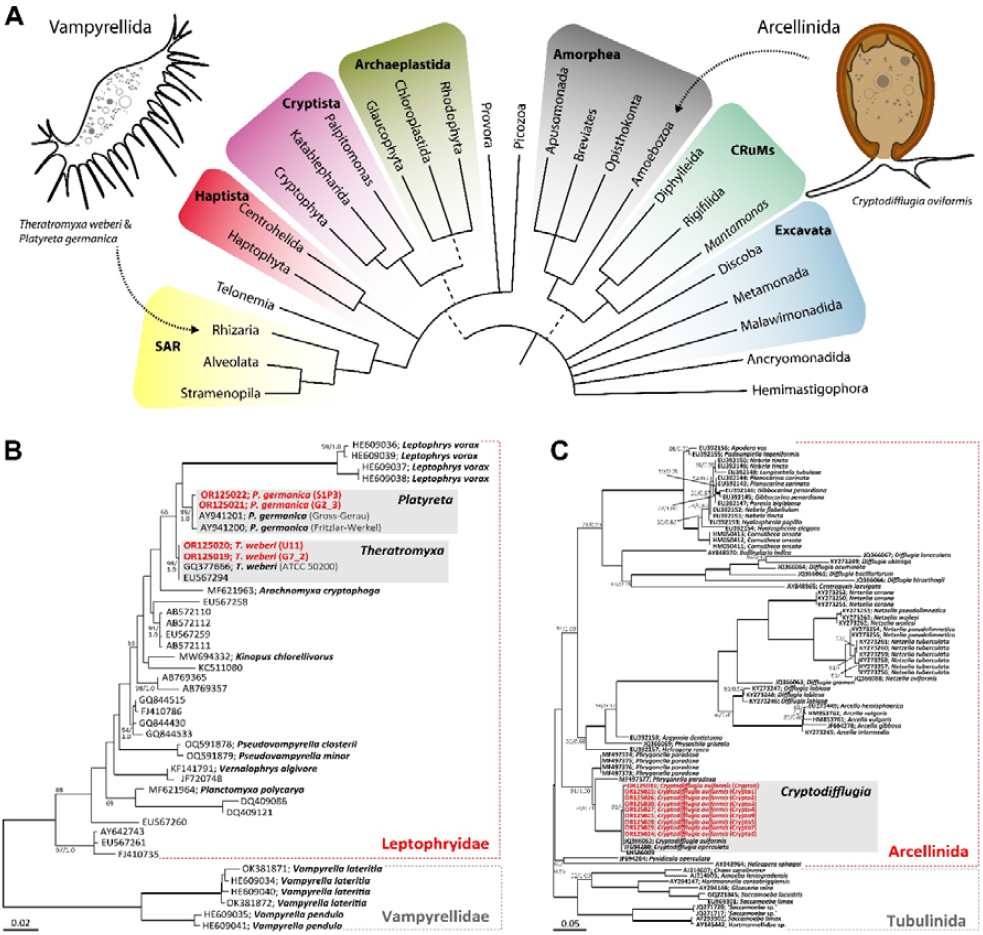
Phylogenetic position of the studied amoebae. (A) Overview of the eukaryotic diversity based on Burki et al. (2020) with highlighted supergroups. Representative drawings indicate the overall morphologies (naked filose vs. testate lobose) and the phylogenetic affinity of the studied amoebae. Note that both taxa are morphologically and genetically vastly different. (B) ML phylogeny of the Leptophryidae (Vampyrellida, Rhizaria) with Vampyrellidae as outgroup. New strains are shown in red. (C) ML phylogeny of the Arcellinida (Amoebozoa) with the Tubulinida as outgroup. New strains are shown in bold. Branches of both phylogenies show bootstrap support and posterior probabilities (ML/BI) unless fully supported (bold) or with support below 50/0.9 (omitted). Branches show bootstrap support and posterior probabilities (ML/BI) unless fully supported (bold) or with support below 50/0.9 (omitted). Scale bars: 0.02 expected substitutions per site

### Morphology and feeding processes of vampyrellid strains

Both vampyrellid species *Theratromyxa weberi* and *Platyreta germanica* had colorless, highly branched cells, best observed with phase contrast optics (Figure 2A, B). The amoeboid cells (= trophozoites) exhibited thin filopodia without granules and moved by slow enlargement and advancement of pseudopodial structures. They showed a high propensity to form network-like structures of unlimited size in our cultures (Figure 2C; Supplementary Video 1). Although both species could be readily grown with yeast as food, they differed markedly in their feeding strategies. *Theratromyxa weberi* incorporated entire yeast cells and cleared the bottom of the culture flask (Figure 2A), while *Platyreta germanica* perforated the yeast cells and phagocytosed the cytoplasm (protoplast feeding). As typical for all known vampyrellids (Hess and Suthaus, 2022), the trophozoites of the studied strains entered an immobile, but highly active digestive cyst stage after feeding (Figure 2D, E). The different feeding strategies observed in *Theratromyxa* and *Platyreta* were also evident from the undigested food remnants left behind in the empty cyst walls. *Theratromyxa* cysts contained an agglomerate of partially digested yeast cells (Figure 2F), while *Platyreta* cysts contained an amorphous, dark brown food remnant (Figure 2G). Such amorphous food remnants can also be found in other protoplast-feeding vampyrellids (Hess et al., 2012), and help identify the mode of feeding before the process itself was observed. As a result of *Platyreta’s* feeding strategy, empty yeast cells accumulate as the cultures develop (Figure 2B). Scanning electron microscopy revealed that these empty yeast cells exhibit large perforations caused by *Platyreta* (Figure 2H).

**Figure 2:**
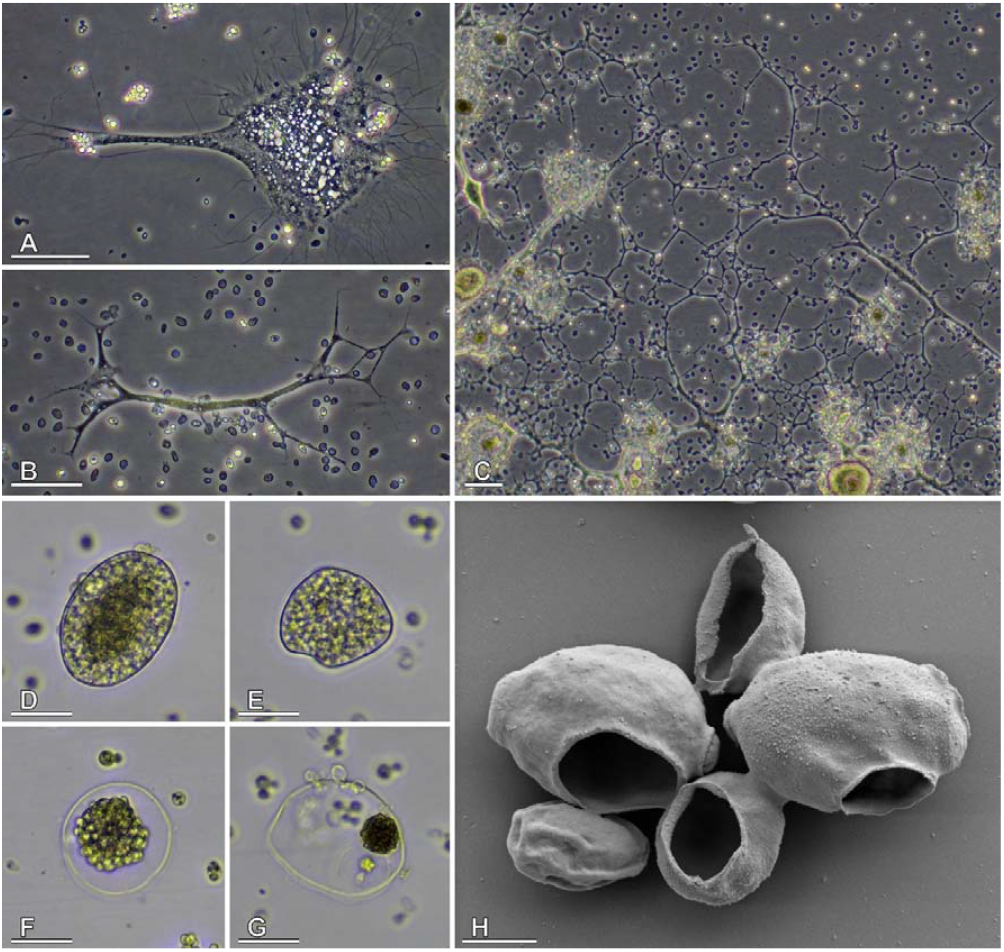
Morphology of soil-dwelling vampyrellids (strains G2.3 and G7.2) and aspects of their feeding behavior. (A) Trophozoite of *Theratromyxa weberi* strain G7.2, phase contrast. (B, C) Trophozoites of *Platyreta germanica* strain G2.3, phase contrast. Note that the cells can fuse and form large networks. (D, E; see also Supplementary Video 1) Full digestive cysts of *T. weberi* (D) and *P. germanica* (E), brightfield. (F, G) Empty digestive cysts left behind by *T. weberi* (F) and *P. germanica* (G) after digestion of the food, brightfield. Note the difference in the food remnants between the engulfing (F) and extracting (G) vampyrellid. (H) Scanning electron micrograph of yeast cells (*Saccharomyces cerevisiae*) perforated and emptied by *P. germanica*. Scale bars indicate 50μm (A-C), 20μm (D-G) and 2μm (H).

### Morphology and feeding processes of Cryptodifflugia strains

*Cryptodifflugia oviformis* cells exhibited an oval, smooth and transparent shell with a mean length of 19 ± 3 μm and a width of 15 ± 3 μm (n=25; measured on strain Crypto3). The opening of the shell was terminal and circular with a diameter of 4 ± 0.6 μm (n=17). These characters and measurements match the current circumscription of the species *C. oviformis* very well (Bobrov and Mazei, 2017). The cells bore several lobose and conical pseudopodia, which reached a length of up to 45 μm. *Cryptodifflugia oviformis* used these pseudopodia for locomotion and to initiate contact with fungal cells. Small fungal structures such as yeast cells and small spores were pulled through the shell’s opening and phagocytosed entirely. We observed that the cells of *Saccharomyces cerevisiae* were frequently deformed during this process, indicating considerable forces applied to the prey cells (Fig. 3E; Supplementary Video 3). If cells were too large to be pulled through the shell’s opening, the fungal cell wall ruptured and the amoebae incorporated both cytoplasm and cell wall material. Relatively thick hyphae were perforated by this process, resulting in a hole with rough edges, while thinner hyphae were broken in two and dragged into the shell (Figure 3A-D; Supplementary Video 2). To study the distribution of F-actin during food uptake, we stained fixed *Cryptodifflugia* cells with a fluorescent phalloidin conjugate. The cell body of the amoebae contained conical bundles of F-actin that started from the point of contact with the fungal cells and extended far into the cell body (Figure 3F). Often these bundles reached the epipodia, i.e. the contact zones of the amoeba’s cell body with the inner wall of the shell (Figure 3F). In motile, foraging amoebae, we did not observe a distinct phalloidin staining in the cell body, but a strong signal in the pseudopodia used for locomotion (Figure 3G).

**Figure 2:**
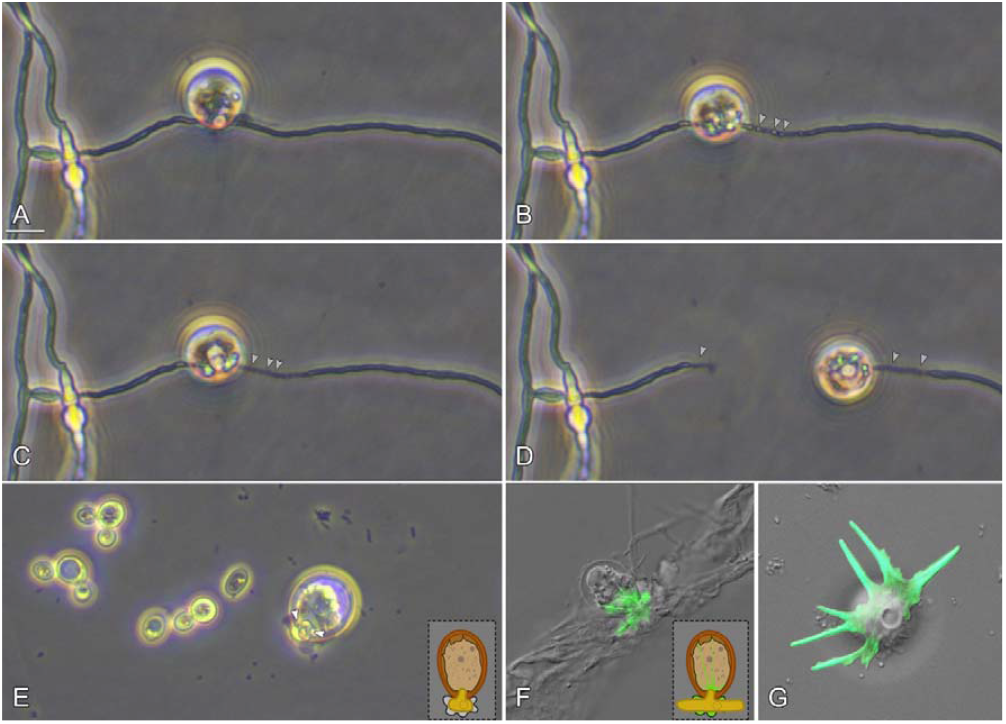
The figure shows the fungivorous protist *Cryptodifflugia oviformis*. Figures A-D show a time series of *C. oviformis* feeding on hyphae *of Fusarium sporotrichoides* (see also Supplementary Video 2). The protist attaches to the hypha and starts to manipulate, pull and break the fungal cells (A). The hypha’s cell wall is broken (B), note that the fungal cell’s contents are fragmented, but still inside of the hypha (see grey arrows). *C. oviformis* empties the fungal hypha (C) and migrates along the hypha while phagocytosis further parts of it (D). Note the grey arrows that indicate empty fragments of the hypha. Figure E shows *C. oviformis* feeding on a yeast cell (*Saccharomyces cerevisiae*). Note that the ovoid and spherical yeast cell is forcefully deformed while being pulled into the shell of the protist (see also drawing and Supplementary Video 3). Figures F-G show F-actin-stained cells. During the rupture of fungal hyphae (here *Amniculicola* sp.), F-actin bundles stretching through the cell body of the protist could be seen (F). During locomotion the cell body was almost entirely devoid of stained F-actin, while pseudopodia were highly stained. Scale bar indicates 10 μm.

### Consumption of fungi by the isolated soil amoebae

Our food range experiments indicated that a variety of fungi can be consumed by the fungivorous protists *Platyreta germanica, Theratromyxa weberi*, and *Cryptodifflugia oviformis*. The range of fungal cultures that was preyed upon by each protistan species differed, as did the mechanisms that were used to access the fungal prey. *Platyreta germanica* perforated the cell walls of its prey and extracted the fungal cell’s protoplast. We did not record a single instance of *Platyreta germanica* phagocytosing entire prey cells. In contrast, the closely related *Theratromyxa weberi* most often phagocytosed entire prey organisms (spores, yeasts, and small hyphae). However, occasional perforation of spores of *Umbelopsis isabellina* and *Mortierella* sp. and the yeasts *Saccharomyces cerevisiae* and *Cryptococcus aerius* were observed as well. *Cryptodifflugia oviformis* consumed small prey cells entirely, but perforated or ruptured hyphae. The hyphae of *Mortierella* sp. and *Rhizopus* sp. showed circular perforations with irregular margins, while we observed all other filamentous fungi to be cut in two during predation. *Cryptodifflugia oviformis* also consumed environmental bacteria, which, however, were not determined.

## 4. Discussion

### Mechanisms of fungivory in amoebae

Fungivory by soil amoebae has been documented for more than 50 years. However, the first reports were still a mystery: Old (1967) observed peculiar, round perforations in the walls of fungal spores which had been incubated in natural soils, but the true nature of the “perforating agent” remained unknown for several years (Old and Wong, 1976; Clough and Patrick, 1976). Finally, Old (1977) and shortly afterward Anderson and Patrick (1980) provided evidence that the perforations were caused by giant terrestrial amoebae. The latter authors also documented several distinct amoebae, which caused perforations of different sizes (from 1 to 5 μm). In our opinion, the applied genus names (e.g. *Vampyrella, Arachnula*) are, however, incorrect. A similar fungivorous amoeba was later cultivated and studied by Pakzad (2003), and finally described as a new taxon, *Platyreta germanica* Bass et al. (2009). It was also shown that this amoeba belongs to the rhizarian protists (and not the Amoebozoa) – relatively closely related to the genus *Vampyrella* (Bass et al., 2009a; Hess et al., 2012). During the past 10 years, the vampyrellid amoebae have been extensively explored by cultivation and molecular methods (Hess et al., 2012; Berney et al., 2013; Gong et al., 2015; Hess, 2017a, 2017b; More et al., 2018, 2021; Zhang et al., 2022), but these studies mainly focused on aquatic representatives. For the terrestrial vampyrellids preying on fungi and animals, we still lack the link between in-depth ecological observations and the genetic identity of particular species. Furthermore, there were no cultures available for experimentation, as all strains studied before were lost. In this study, we isolated *Platyreta germanica* from the rhizosphere of *A. thaliana* and established a bacteria-free culture for our feeding experiments. In addition, we demonstrate that two other terrestrial amoebae, *Theratromyxa weberi* and *Cryptodifflugia oviformis* readily feed on diverse fungi – including hyphal structures. This came as a surprise, as *T. weberi* was originally described as a carnivorous amoeba feeding on nematodes (Weber et al., 1952; Sayre, 1973). It was also shown that a *Cryptodifflugia* species is able to consume nematodes (Geisen et al. 2015). Hence, we presume that several soil amoebae have a fairly wide prey range.

Furthermore, our results exemplify that fungivory is not a uniform protistan trait. All three studied soil amoebae, *Platyreta germanica, Theratromyxa weberi* and *Cryptodifflugia oviformis*, were able to manipulate, ingest and/or perforate fungal cells. However, how exactly these amoebae fed on the same fungal prey differed strikingly – even among the two very closely related vampyrellid species. While *Platyreta* extracts fungal cells after local perforation of the wall, *Theratromyxa* usually ingested entire cells, which might have an impact on the possible prey range (size-limited ingestion). In fact, *Theratromyxa* showed a somewhat narrower prey range and was not able to feed on the filamentous *Fusarium* species. The different prey ranges of *Platyreta germanica* and *Cryptodifflugia oviformis* cannot be explained by prey size, as both organisms can access the cytoplasm of spores and hyphae by the destruction of the fungal wall. How exactly these amoebae open fungal cells remains unclear. Based on the regular morphology and smooth margins of the perforations produced by vampyrellid amoebae such as *Platyreta*, the action of enzymes was hypothesized (Old, 1978) but not tested. In fact, the annular perforations of *Platyreta* closely resemble those found in other rhizarian protoplast feeders (*Orciraptor agilis*) which evidently use lytic carbohydrate-active enzymes to open prey cell walls (Moye et al., 2022). Besides *Cryptodifflugia*, there are a few reports of other arcellinid amoebae preying on and/or perforating large organisms. These observations were mainly made on limnic species (Hoogenraad and Groot, 1941; Siemensma and Opitz, 2014; Geisen et al., 2015b; Dumack et al., 2018a). For example, *Difflugia rubescens* and *Pseudonebela africana* were found to perforate cell walls of green algae, leaving roughly-edged holes in the remaining cell wall (Hoogenraad & Groot, 1941; Siemensma & Opitz, 2014). Dumack et al. (2018) speculated that this process is mechanical, potentially being facilitated by the cytoskeleton of these amoebae. They observed *Phryganella paradoxa*, a specialized predator of diatoms and close relative of *Cryptodifflugia oviformis* (see Figure 1C), to bend and break the siliceous thecae of their prey. The aperture of the amoeba’s shell seemed to function as a pivot point in this process and thus its diameter might be the limiting factor in terms of prey size. Here, we found F-actin bundles in attacking cells of *Cryptodifflugia oviformis*, which supports the idea that the cytoskeleton in testate amoebae exerts mechanical force during the feeding process. In contrast to the diatom-feeding *Phryganella paradoxa*, the prey range of *Cryptodifflugia oviformis* seems not to be limited by prey size, which we explain by the flexibility of fungal cells (e.g. note the deformation of yeasts cells when passing the aperture). Hence, the observed differences in prey range between *Cryptodifflugia oviformis* and the vampyrellid amoebae may also be related to the evolutionary and functionally different mechanisms of fungivory in these two distantly related groups of amoebae. We conclude that fungivory evolved repeatedly in protist lineages, and that fungivorous soil amoebae evolved functionally distinct strategies that result in different prey ranges. Fungivory in protists is thus not a uniform trait, and depending on whether the feeding pressure is directed at yeasts, fungal spores, filamentous fungi or all of them, fungivorous protists might have different functions in the soil food web.

### Fungivory and its consequences for the soil food web

*Arabidopsis thaliana* is widely used as a model plant, but associated eukaryotic microorganisms remain largely unexplored (Urbina et al., 2018). Although *A. thaliana* does not engage in mycorrhizal symbiosis, the growth and health of *A. thaliana* depend on root-associated fungi (Micallef et al., 2009; Urbina et al., 2018). Junker et al. (2012) isolated fungi from *Arabidopsis thaliana*. Among others, they found *Fusarium* sp. and *Rhizopus* sp., species that – as we demonstrated in our experiments – were consumed by *Platyreta germanica* (*Fusarium* spp.) and *Cryptodifflugia oviformis* (*Fusarium* spp. and *Rhizopus* sp.). Yeasts are neglected in rhizosphere studies, but evidence by stable isotope probing suggests that yeasts could be important competitors of rhizobacteria for root exudates and there are numerous protist species feeding on yeasts (Dumack et al., 2018b; Hünninghaus et al., 2019). Geisen et al. (2015) showed that vampyrellid amoebae represent a small fraction of the protist community, but they are consistently present in various terrestrial habitats (see also Fiore-Donno et al., 2019, 2020). *Cryptodifflugia oviformis* is long known to be a ubiquitous and abundant protist (Bobrov and Mazei, 2017), which we found along the roots of *Arabidopsis thaliana* in high numbers. The amount of energy that is channeled by fungivorous protists may thus be considerable (Hedley et al., 1950; Kramer et al., 2016). Here, we provide publicly available cultures of three potentially important soil amoebae, which may facilitate future experimental research to understand the functions, relevance and potential use of fungivorous protists.

## Acknowledgments

This work was funded by the German Research Foundation in the framework of the priority programme SPP 1991 (Taxon-Omics: New approaches for discovering and naming biodiversity), grant 447190101 to S.H. and 447013012 to M.B. and K.D.

Figures and tables; may be printed in black and white.

## Notes

### Competing Interest Statement

The authors have declared no competing interest.

